# Shared Alteration of Whole-Brain Connectivity and Olfactory Deficits in Multiple Autism Mouse Models

**DOI:** 10.1101/2025.02.02.636175

**Authors:** Tsan-Ting Hsu, Chih-Ping Chen, Ming-Hui Lin, Tzyy-Nan Haung, Chung-Yu Wang, Chia-Ming Lee, Bi-Chang Chen, Chien-Yao Wang, Yi-Ping Hsueh

**Author notes:** Correspondence to: Yi-Ping Hsueh, Institute of Molecular Biology, Academia Sinica,; Chien-Yao Wang, Institute of Information Science, Academia Sinica,; Bi-Chang Chen, Research Center for Applied Sciences, Academia Sinica,. Mailing address: Academia Sinica, 128, Sec. 2, Academia Rd., Taipei, 11529, Taiwan, ROC. Authors’ email addresses.

## Abstract

Autism spectrum disorder (ASD) is a disconnection condition influenced by both heterogeneous genetic and environmental factors, yet it remains unclear whether common connectivity deficits exist. Here, we demonstrate that different ASD-linked mutations lead to distinct circuit abnormalities but share deficits in the piriform cortex and olfactory discrimination. Using advanced artificial intelligence, we developed a whole-brain mapping platform to analyze the distribution of the Thy1-YFP projection neurons in three ASD mouse models (*Tbr1^+/–^*, *Nf1^+/–^*, *Vcp^+/R95G^*). Our analysis revealed changes in axonal patterns and neuronal distribution, indicating deficits in projection neuron differentiation and maintenance. Notably, the piriform cortex consistently exhibited reduced YFP^+^ cells and signals and impaired functionality across all models. Visual and somatosensory cortices were also affected, but the patterns varied. These findings highlight that the sensory regions, especially the piriform cortex, are susceptible to ASD-related mutations, strengthening the notion that different sensory experiences are common in ASD.

**Highlights:**

- AI-powered whole-mouse brain quantification accelerates neural connectivity study.
- Autism-linked mutations lead to various circuit abnormalities of Thy1-YFP neurons.
- Multiple sensory regions all exhibit circuit deficits among the three autism models.
- Abnormalities of olfaction and piriform cortex circuits are common in autism models.

## Introduction

Autism spectrum disorders (ASD) are highly heterogeneous neuropsychiatric conditions influenced by diverse genetic and environmental factors during development ^1–3^. Despite the variability in behavioral symptoms, ASD is characterized by two core features: deficits in social interaction and verbal/nonverbal communication, along with repetitive behaviors and sensory abnormalities ^4^. Abnormal synaptic connectivity, encompassing both hypo-and hyper-connectivity, is a hallmark of ASD ^5–8^. Given the involvement of multiple brain regions in ASD, identifying common connectivity deficits across different ASD conditions remains a significant challenge. If such shared deficits exist, they are likely to play critical roles in ASD etiology.

Over the past decades, structural and functional magnetic resonance imaging (sMRI and fMRI) have been widely used to analyze brain abnormalities in ASD patients ^9–12^ and in rodent models with ASD-associated mutations ^13–19^. Rodent models, with their relatively homogeneous genetic backgrounds and controlled environmental factors, provide valuable insights into how specific genetic variations associated with ASD influence brain morphology and functional connectivity ^16,18^. MRI studies have revealed features of ASD such as brain volume enlargement in young children ^9,20^, defects in the mesolimbic reward system ^10^, disrupted interhemispheric connectivity ^21^, and hyperfunctional connectivity caused by Tsc2 haploinsufficiency ^16^.

Beyond macroscale connectivity analysis via MRI, mesoscale connectivity analysis using whole-brain immunostaining and registration to a standard mouse brain has also been developed ^22,23^. This approach enables detailed analysis of projections from specific cell types and subregions (https://connectivity.brain-map.org), further providing insights into circuit deficits caused by ASD conditions. Recently, we established a whole-brain immunostaining and quantification platform to study the anterior basolateral amygdala (BLAa) projections in *Tbr1*^+/−^ mice ^24^, a model of *TBR1* haploinsufficiency in ASD ^25–28^. Using the channelrhodopsin variant oChIEF fused with Citrine, we visualized BLAa axonal projections and analyzed functional connectivity of BLAa based on *c-Fos* expression in various brain regions ^24^. Our findings indicate that *Tbr1* haploinsufficiency disrupts contralateral BLAa projections, alters global BLAa connectivity, and impairs whole-brain synchronization under both resting and theta-burst stimulation (TBS) at BLAa conditions. Notably, TBS-induced activation at the BLAa also enhanced social interactions in *Tbr1*^+/−^ mice ^24^. These results reveal dysconnectivity and abnormal synchronization of the BLAa in an ASD model and suggest a potential mechanism for improving social behaviors through targeted stimulation.

While studying specific regions, such as the BLAa, provides valuable insights into ASD-related connectivity and function, analyzing individual brain areas is labor-intensive. To address this challenge, we here enhanced the accuracy and efficiency of our whole-brain mapping and quantification platform by integrating advanced artificial intelligence for image registration, specifically DeepReg. This AI tool allowed us to accelerate the brain mapping and quantification process with automated region-of-interest (auto-ROI) correction. We thus named this system BM- auto (brain mapping with auto-ROI correction). Using BM-auto, we analyzed Thy1-YFP reporter expression in three mouse models with ASD-associated mutations (*Tbr1*^+/−^, *Nf1*^+/−^, and *Vcp*^+/*R95G*^) ^27,29,30^. In Thy1-YFP-H transgenic mice, YFP highlights axons and specific projection neurons in the hippocampus, cortex, and amygdala ^31^, making it a valuable tool for studying altered connectivity patterns in ASD models.

Mutations in *Tbr1*, *Nf1*, and *Vcp* result in social and synaptic deficits despite their distinct molecular functions and expression patterns ^27,29,30,32^. *Tbr1* encodes a neuron-specific transcription factor critical for the development of projection neurons in the cortex and amygdala ^27,33–35^. As mentioned above, we showed that TBR1 controls both the structural and functional connectivity of BLAa in mice. Moreover, the deficits of the posterior part of the anterior commissure, a white matter structure linking two BLAa in the two brain hemispheres, share the evolutionary conserved deficits in humans and mice with TBR1 deficiency ^36,37^. It is certainly intriguing to investigate if *Tbr1* haploinsufficiency influences the connectivity of other brain regions, particularly the cerebral cortex.

*Nf1* encodes a scaffold protein controlling the Ras and cAMP pathways, as well as endoplasmic reticulum (ER) formation via the interaction with valosin-containing protein (VCP) ^29,38^. *Vcp* encodes an ATPase chaperone involved in multiple cellular processes, including ER formation ^39,40^. Our previous studies showed that NF1 protein and VCP interact to control ER formation and consequent protein synthesis, ultimately dendritic spine formation ^29,30,40^. Although both *NF1* and *VCP* are associated with ASD ^41–45^, mutation of these two genes also results in other neurological diseases ^41,42,46^. As far, it is unclear if *Nf1* and *Vcp* share the same deficits in neural circuits. Together with *Tbr1*, we anticipated that if the mutations of these three genes result in common connection defects, those deficits are likely critical to ASD etiology.

Based on these assumptions, we here use our BM-auto platform to analyze the Thy1-YFP reporter in *Tbr1^+/–^*, *Nf1^+/–^* and *Vcp^+/R95G^* mice. We found that YFP signal highlighted various axonal innervation patterns. Thy1-YFP cell numbers also changed in different brain regions, revealing deficits in the differentiation or maintenance of projection neurons. Altered neural connections were predicted based on the Allen Mouse Brain Connectivity Atlas and our data. Among all comparisons, the olfaction-associated piriform cortex proved to be the only region displaying common defects in all three mutant mice. Consistently, we detected common impairments in olfactory discrimination. Visual and somatosensory cortices were also affected in the mutant mice, although the tendencies differed. Overall, our whole-brain analysis demonstrates that the connections and functions of sensory-related brain regions are susceptible to various ASD-related dysfunctions, echoing abnormal sensations in ASD.

## Results

### Whole-brain mesoscale imaging

We have established two sets of whole-brain imaging systems. One is light-sheet microscopy combined with a PEGASOS clearing method to acquire continuous coronal mouse brain images (**Figure 1A**) ^47,48^. The other is high-content imaging of serial coronal mouse brain sections (**Figure 1B**) ^24^. The horizontal view of 3D images confirmed the dominant expression of Thy1-YFP in the hippocampal formation (HPF), basolateral amygdala (BLA), and cerebral cortex (**Figure 1B, 1B Supplemental Video 1, 2**). We noticed that YFP signals or YFP^+^ cell numbers in some cortical regions of our images, including the primary somatosensory cortex (SSp) and primary visual cortex (VISp), were higher in *Tbr1^+/–^* mice, a mouse model of ASD established previously, compared to wild-type (WT) littermates (**Figure 1A, 1B**), prompting subsequent quantification. Although light-sheet microscopy revealed a high-quality 3D continuous brain map (**Figure 1A, Supplemental Video 1**), high-content imaging of serial brain sections resulted in a better resolution of axonal projections to individual coronal sections (**Figure 1C, Supplemental Video 1** vs. **2**). Therefore, we used the results of high-content imaging for further quantification analysis.

**Figure 1.**
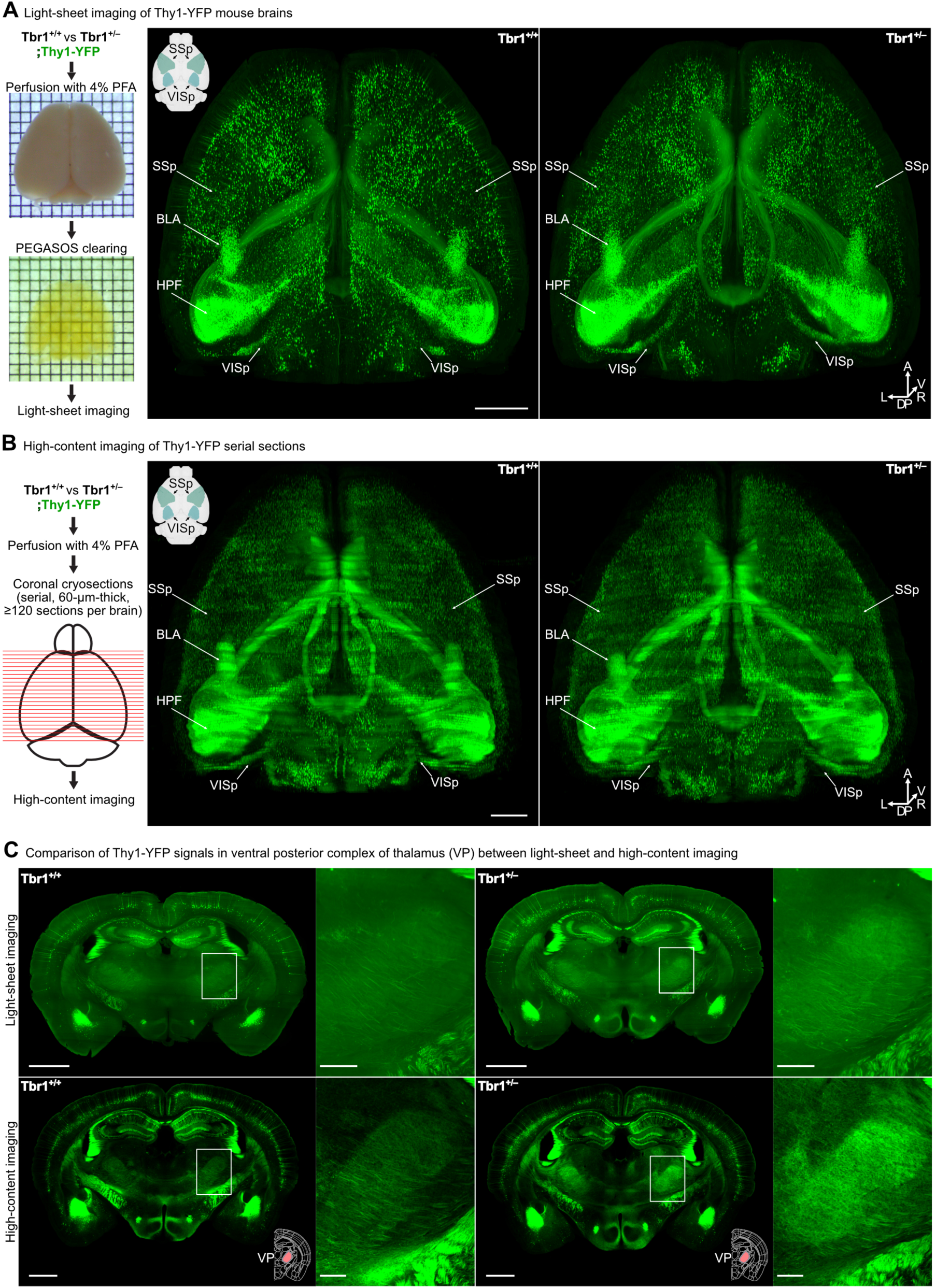
Imaging of *Tbr1*^+/+^;Thy1-YFP and *Tbr1*^+/–^;Thy1-YFP mouse brains by two methods. (**A**) Light-sheet imaging of *Tbr1*^+/+^;Thy1-YFP and *Tbr1*^+/–^;Thy1-YFP mouse brains. The left panel illustrates the workflow comprising PFA fixation, PEGASOS clearing, and light-sheet imaging. Grids in the photo represent 1 mm^2^. The panel at right shows 3D projection–horizontal views of the *Tbr1*^+/+^;Thy1-YFP and *Tbr1*^+/–^;Thy1-YFP mouse brain (voxel size: LR: 10 µm x DV: 10 µm x AP: 10 µm). Arrows indicate the SSp and VISp exhibiting differential Thy1-YFP signals between genotypes. The areas of the brain regions are illustrated in the upper left insert. (**B**) High-content imaging of Thy1-YFP coronal brain slices. The left panel illustrates the workflow encompassing PFA fixation, cryosectioning into coronal slices, and high-content imaging. The panel at right shows 3D projection–horizontal views of the *Tbr1*^+/+^;Thy1-YFP and *Tbr1*^+/–^;Thy1-YFP image stacks (voxel size: LR: 10 µm x DV: 10 µm x AP: 60 µm). The SSp and VISp, which show differential Thy1-YFP signals between genotypes, are also indicated. The insert in the upper left corner illustrates the areas of brain regions. (**C**) Comparison of Thy1-YFP signals between light-sheet and high-content imaging. The upper panel shows image stacks (60 µm-thick) of six coronal optical sections acquired by light-sheet imaging of the *Tbr1*^+/+^ (upper left) and *Tbr1*^+/–^ (upper right) mouse brain. The lower panel shows single coronal sections (60 µm-thick) acquired by high-content imaging of the *Tbr1*^+/+^ (lower left) and *Tbr1*^+/–^ (lower right) mouse brain with similar AP coordinates as the image stacks of light-sheet imaging. Boxed areas show Thy1-YFP signals in the VP (illustrated in the bottom right panels) and enlarged views are shown next to the whole-slice images. Scale bar of Thy1-YFP images in (**A**-**C**), 1 mm; boxed area in **c**, 200 μm.

### Whole-brain mapping and quantification with automatic correction (BM-auto)

We modified our recently established whole-brain registration and quantification platform ^24^ to analyze the differential distribution of both YFP signals and YFP^+^ cell numbers in the *Tbr1^+/–^*, *Nf1^+/–^* and *Vcp^+/R95G^* mouse brains (**Figure 2**). We used WT littermates of mutant mice, denoted *Tbr1^+/+^*, *Nf1^+/+^* and *Vcp^+/+^* respectively, for comparison. We took a similar sample size for analyses based on previous histology-based whole-brain studies ^24,49,50^. Each group comprised six mutant mice and six WT individuals. To ensure accurate quantification, first, we manually corrected the region of interest (ROI) for 37 brain regions in all three sets of comparisons. These 37 regions exhibited differential YFP signals or YFP^+^ cell numbers for one or multiple comparisons. Furthermore, using these 37 ROI-corrected masks and data on background tissue autofluorescence, we established an automatic ROI correction platform (auto-ROI correction) to perform deep learning-based registration (DeepReg) (**Figure 2**). Our auto-ROI correction deploying DeepReg estimated dense deformation fields from the Allen Mouse Common Coordinate Framework version 3 (CCFv3) template applied to our original brain images. The predicted deformation fields were then applied to the CCFv3 region masks, resulting in auto-ROI-corrected masks, which were then applied to the fluorescent images for quantification (**Figure 2**, see Methods for details). Our proposed DeepReg model achieved substantial improvements over current state-of-the-art methods. Consequently, our auto-ROI correction effectively minimized misalignments between region mask boundaries and the brain image, resulting in enhanced quantification accuracy. We name the entire platform as BM-auto. Details of the performance of different registration methods are summarized in **Table 1**.

**Figure. 2.**
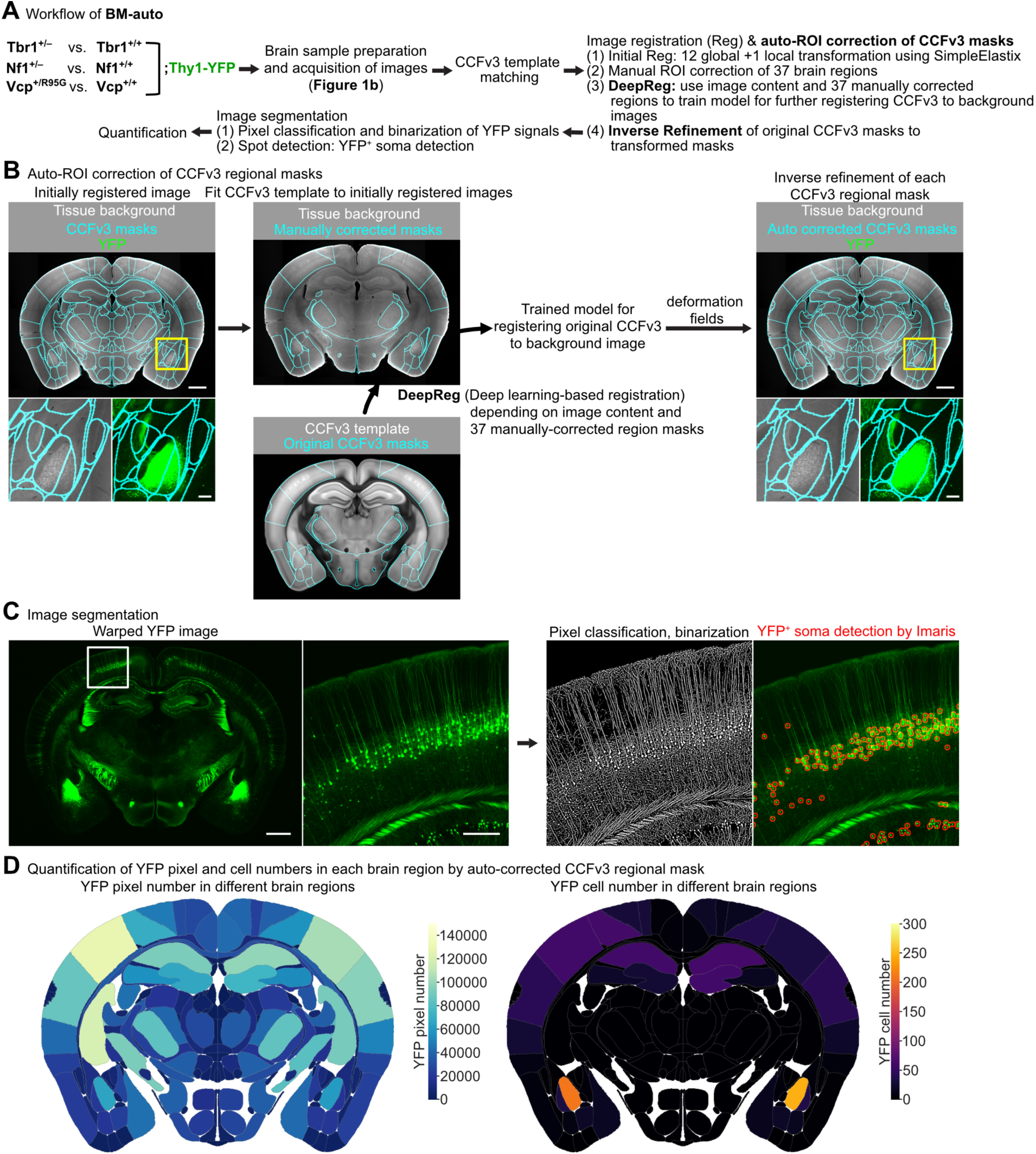
Workflow of whole-brain analysis of Thy1-YFP signals. (**A**) The flowchart of BM-auto, including all types of brain samples used for high-content imaging, subsequent image registration, together with auto-ROI correction of Allen Mouse Common Coordinate Framework version 3 (CCFv3) regional masks, image segmentation, and quantification. (**B**) The workflow of auto-ROI correction of CCFv3 regional masks. The left panel shows the unsuitable alignment of the original CCFv3 regional masks with the initially registered image. Consequently, we developed auto-ROI correction of CCFv3 regional masks. The middle to right panels show the procedures for developing auto-ROI correction. CCFv3 templates were trained to fit initially registered images with tissue background by image content and 37 manually ROI-corrected regional masks (manually-corrected ROIs, middle) using deep learning-based registration (DeepReg). The trained DeepReg model was applied to register each CCFv3 template to the corresponding background image. The resulting deformation fields were used to inversely refine the original CCFv3 regional masks to auto-corrected CCFv3 regional masks (auto-corrected ROIs). (**C**) The YFP signals and soma locations of Thy1-YFP neurons were segmented and detected respectively from the initially registered Thy1-YFP images (also see **Supplemental Figure 3). (D**) Auto-corrected and 37 manually-corrected ROIs were applied to quantify YFP signals (pixel numbers, choropleth in left panel) and YFP^+^ cell numbers (choropleth in right panel) of different brain regions in a single coronal section. Scale bar of tissue background and Thy1-YFP images in (**B, C**), 1 mm; boxed area in (**B**), 200 μm; boxed area in (**C**), 300 μm.

**Table 1.** Performance details of the different registration methods.

| Method | Dice ↑ | Boundary Dice ↑ | HD95 (pixels) ↓ | Negative Jacobian (%) ↓ |
| --- | --- | --- | --- | --- |
| Manual Matching | 0.3479 | 0.1248 | 54.1534 | - |
| Affine Registration | 0.5732 | 0.3269 | 26.8697 | 0 |
| **Elastix (Initial Registration)** | **0.7190** | **0.5522** | **13.2960** | **0.0227** |
| **Deep learning-based methods** |  |  |  |  |
| VoxelMorph <sup>85</sup> | 0.8066 | 0.6102 | 11.6875 | 0.4619 |
| TransMorph <sup>86</sup> | 0.8153 | 0.6348 | 11.0724 | 0.4076 |
| VMambaMorph <sup>87</sup> | 0.8049 | 0.6206 | 11.2939 | 0.2627 |
| ModeT <sup>88</sup> | 0.7953 | 0.5921 | 12.2801 | 0.0479 |
| VFA <sup>89</sup> | 0.8316 | 0.6400 | 11.3467 | 0.0151 |
| CorrMLP <sup>82</sup> | 0.8297 | 0.6494 | 10.6373 | 0.9688 |
| FourierNet <sup>83</sup> | 0.7912 | 0.5929 | 12.1432 | 0.2487 |
| **DeepReg (this study)** | **0.8512** | **0.7049** | **9.0756** | **0.1923** |

1. ↑ : higher value is better. ↓ : lower value is better. 2. Dice: the Dice similarity coefficient <sup>90</sup>. 3. Boundary Dice: the Boundary Dice similarity coefficient <sup>51</sup>. 4. HD95 (pixels): the 95th percentile Hausdorff distance <sup>91</sup>. 5. Negative Jacobian (%): the percentage of pixels with a negative Jacobian value <sup>92</sup>.

The effectiveness of the auto-ROI corrections was first assessed by comparing Boundary Dice scores ^51^ with or without auto-ROI corrections (**Supplemental Figure 1A**). Boundary Dice scores evaluate the accuracy of boundary pixel alignment between predicted and ground truth regions. Our results show that the auto-ROI corrections significantly reduced registration error in the subcortical regions and fiber tracts (fb) of the coronal brain slices compared to the initial registration. Overall, the Boundary Dice scores for subcortical regions and fb improved by 30% to 600%, which proved crucial for subsequent quantitative analysis (**Supplemental Figure 1A**). Next, we overlapped the auto-corrected region masks with manually-corrected region masks and found that they largely matched (**Supplemental Figure 1B**), indicating good auto-ROI correction performance by our DeepReg inverse refinement of CCFv3 region masks. Furthermore, we compared the three datasets, i.e., quantification results for manual, auto and no ROI corrections. The differences in metrics (ΔYFP signal or ΔYFP^+^ cell number (mutant - WT), see Methods) between the auto vs. manual ROI correction and no ROI correction vs. manual ROI correction groups were measured and then compared. In general, the differences in ΔYFP signal or ΔYFP^+^ cell number (i.e., ΔΔYFP or ΔΔYFP cell, respectively) were reduced in many brain regions for the auto vs. manual ROI comparison relative to the no ROI correction vs. manual ROI comparison (**Supplemental Figure 1C, 1D**). Furthermore, we observed better delineation of regional boundaries upon auto-ROI correction (**Supplemental Figure 2**), which increased registration accuracy. These results support the reliability and accuracy of our whole-brain quantification analysis using auto-ROI correction.

After removing regions with very low or no YFP signals, we subjected both hemispheres of more than 300 brain regions, encompassing both grey and white matter, to further analyses (**Supplemental Tables 1-3**). All abbreviations and full names of the brain regions are available on the Allen Brain Reference Atlas website (https://atlas.brain-map.org) and in **Supplemental Tables 1-3**. The YFP signals and Thy1-YFP cell numbers at a whole-brain scale and the potential neuronal circuit deficits caused by *Tbr1*, *Nf1* and/or *Vcp* deficiency were then analyzed based on the altered YFP patterns and Allen Mouse Brain Connectivity Atlas (http://connectivity.brain-map.org).

### Alteration of whole-brain projections

Our BM-auto analysis showed that *Tbr1^+/–^* mice tended to have more brain regions with higher YFP signals than their WT littermates (**Figure 3A, 3B, Supplemental Table 1**; higher in *Tbr1^+/–^*: left hemisphere 101 regions, right 109 regions; lower in *Tbr1^+/–^*: left hemisphere 22 regions, right 10 regions). In contrast, *Nf1^+/–^* mice had more brain regions with lower YFP signals than their WT littermates (**Figure 3C, 3D, Supplemental Table 2**; higher in *Nf1^+/–^*: left hemisphere 14 regions, right 20 regions; lower in *Nf1^+/–^*: left hemisphere 67 regions, right 54 regions). *Vcp^+/R95G^* mice displayed an overwhelmingly dominant reduction in YFP signals, with 254 and 228 brain regions in the left and right hemispheres, respectively, having lower YFP signals. In fact, only three regions in the left and right hemispheres of *Vcp^+/R95G^* mouse brains exhibited higher YFP signals than WT littermates (**Figure 3E, 3F, Supplemental Table 3**). Thus, *Tbr1*, *Nf1* and *Vcp* deficiencies have divergent effects on the distribution of Thy1-YFP in mouse brains.

**Figure 3.**
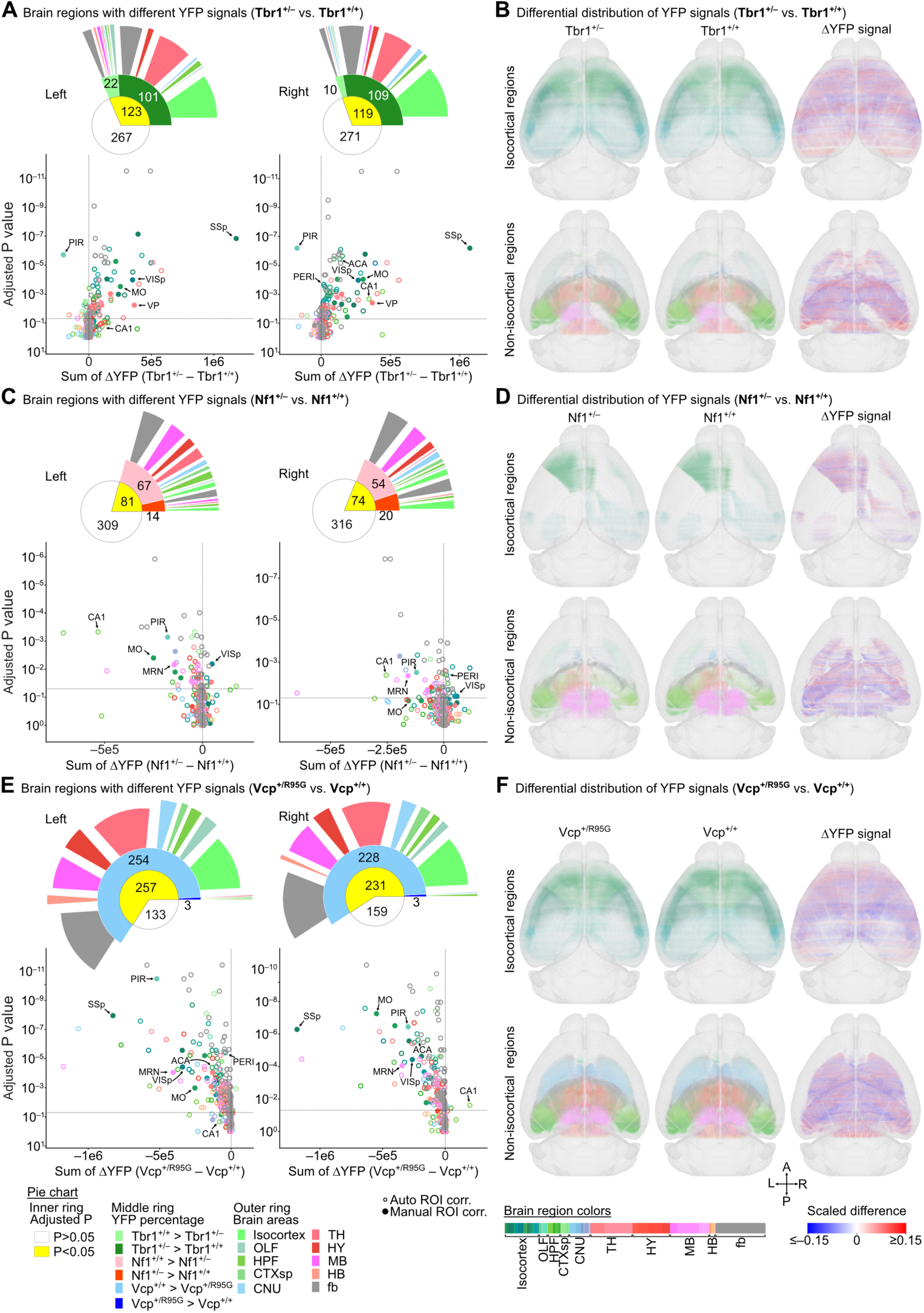
*Tbr1*, *Nf1* and *Vcp* deficiencies result in differential distribution patterns of YFP in Thy1-YFP mouse brains. (**A, C, E**) Region-wise comparison of YFP signal distributions along the anterior–posterior (AP) axis in both left and right brain hemispheres for the (**A**) *Tbr1*^+/–^ vs. *Tbr1*^+/+^, (**C**) *Nf1*^+/–^ vs. *Nf1*^+/+^, (**E**) *Vcp*^+/R95G^ vs. *Vcp*^+/+^ groups. Each scatterplot shows the summed difference of YFP signals along the AP axis (Sum of ΔYFP) versus statistical significance (Adjusted *P* value) for each brain region of the compared groups (also see **Supplemental Tables 1-3**). Friedman’s one-way repeated measures were used for statistical analysis. *P* values were further adjusted to control for the false discovery rate. Vertical and horizontal dotted lines represent the Sum of ΔYFP = 0 and Adjusted *P* value = 0.05, respectively. Colors represent brain region identities (indicated below panel **f**). Arrows point out certain brain regions having differential Thy1-YFP signals in all three compared groups or in specific groups (such as the VP in *Tbr1*^+/–^ vs. *Tbr1*^+/+^, and the SSp in *Tbr1*^+/–^ vs. *Tbr1*^+/+^ and *Vcp*^+/R95G^ vs. *Vcp*^+/+^). Open and filled circles represent auto-ROI-corrected and manual-ROI-corrected brain regions, respectively. The pie charts illustrate the proportions of brain regions that fit the condition annotated at the bottom of panel **E**. (**B, D, F**) Horizontal views of 3D maps with binarized and binned (10 μm^3^) YFP voxels in the brain regions displaying a significant difference (Adjusted *P* value < 0.05) in the left and right hemispheres of the (**B**) *Tbr1*^+/–^ vs. *Tbr1*^+/+^, (**D**) *Nf1*^+/–^ vs. *Nf1*^+/+^, and (**F**) *Vcp*^+/R95G^ vs. *Vcp*^+/+^ groups. As indicated, the left and middle panels are the 3D maps of mutant and WT mouse brains. Colors of YFP voxels in the left and middle panels represent brain regions as shown below (**F**), and the alpha levels (0–1) of the brain region color represent scaled YFP probabilities. The panels at right show the differences in YFP signal probabilities in every 10 μm^3^ voxel (ΔYFP from mutant – WT). The heatmap scale for ΔYFP is shown in the bottom right below panel **F**. A, anterior; P, posterior; L, left; R, right. Sample sizes of mice are identical in (**A**-**F**). A total of six groups of six mice each were analyzed.

In addition to total YFP signals, we also analyzed Thy1-YFP neuron numbers based on the signals in the soma. In general, *Tbr1*, *Nf1* and *Vcp* deficiencies also influenced the numbers of Thy1-YFP neurons in different brain regions (**Figure 4, Supplemental Figure 4**). Similar to our results for total YFP signals, we found that *Tbr1^+/–^* mouse brains had more regions possessing more YFP^+^ cells (**Figure 4A, 4B, Supplemental Table 1**). In contrast to the respective results for YFP signals, *Nf1^+/–^* mice had more regions with more YFP^+^ cells (**Figure 4C, 4D, Supplemental Table 2**). *Vcp^+/R95G^* mice again presented more brain regions having reduced numbers of YFP^+^ cells (**Figure 4E, 4F, Supplemental Table 3**), although there were far fewer affected brain regions than determined for total YFP signals. Thus, the impact of *Tbr1*, *Nf1* and *Vcp* deficiencies on Thy1-YFP neuron numbers in brain regions also varies (**Figure 4**).

**Figure 4.**
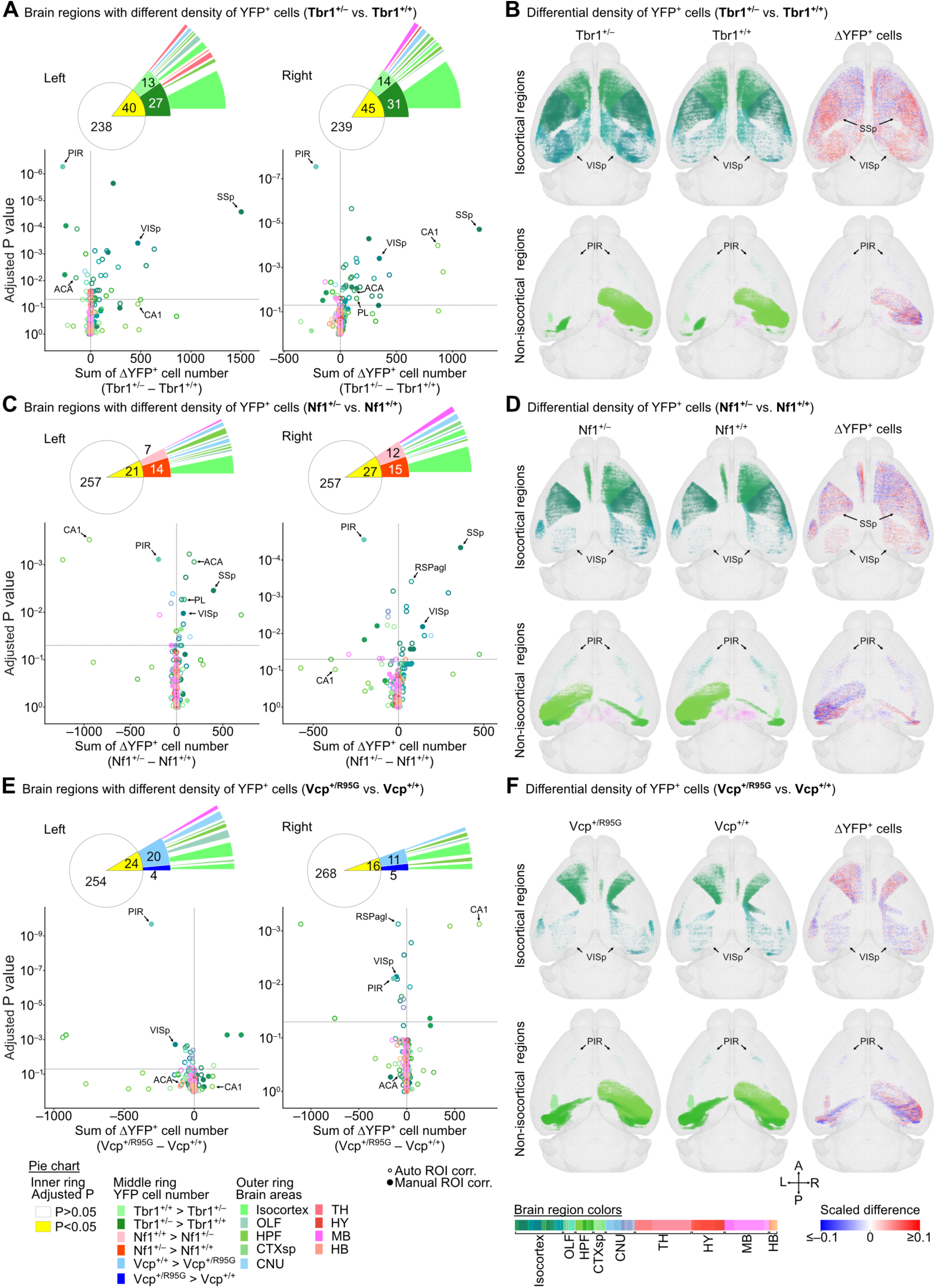
*Tbr1*, *Nf1* and *Vcp* deficiencies differentially alter cell numbers of Thy1-YFP neurons in various brain regions. The same sets of mice analyzed in Figure 3 were used to analyze the numbers of Thy1-YFP neurons at a whole-brain scale. The analyses were conducted in a similar fashion to those described in Figure 3, except that numbers of Thy1-YFP neurons were investigated. (**A, C, E**) Region-wise comparison of YFP^+^ cell distributions along the AP axis in both the left and right brain hemispheres of the (**A**) *Tbr1*^+/–^ vs. *Tbr1*^+/+^, (**C**) *Nf1*^+/–^ vs. *Nf1*^+/+^, and (**E**) *Vcp*^+/R95G^ vs. *Vcp*^+/+^ groups. Each scatterplot shows the sum of ΔYFP^+^ cells along the AP axis versus statistical significance for each brain region of the compared groups (also see **Supplemental Tables 1-3**). Friedman’s one-way repeated measures were used for statistical analysis. *P* values were further adjusted to control for the false discovery rate. Open and filled circles represent auto-ROI-corrected and manual-ROI-corrected brain regions, respectively. The pie charts illustrate the proportions of brain regions that fit the condition annotated below (**E**). (**B, D, F**) Horizontal views of 3D maps with binned (100 μm^3^) YFP^+^ cell cubes if the cubes in the brain regions display a significant difference (Adjusted *P* value <0.05) in the left and right hemispheres of the compared groups: (**B**) *Tbr1*^+/–^ vs. *Tbr1*^+/+^, (**D**) *Nf1*^+/–^ vs. *Nf1*^+/+^, (**F**) *Vcp*^+/R95G^ vs. *Vcp*^+/+^. As indicated, the left and middle panels are 3D maps of the mutant and WT mouse brains. Colors of YFP^+^ cell cubes in the left and middle panels represent brain regions, as shown below (**F**), with their sizes representing the relative normalized density of YFP^+^ cell numbers within the binned cubes. The panels at right show the differences in the density of YFP^+^ cell numbers (ΔYFP^+^ cell number from mutant – WT). The heatmap scale for ΔYFP^+^ cell number is shown at right below (**F**). A, anterior; P, posterior; L, left; R, right. Sample sizes of mice are identical in (**A**-**F**). A total of six groups of six mice each were analyzed.

### Retrograde labeling of the ventral posterior nucleus

Next, we performed retrograde labeling to validate the relevance of the altered YFP signals for differing axonal innervation. Several thalamus nuclei, such as the ventral posterior nucleus (VP), do not possess Thy1-YFP neurons, but did present higher YFP signals in the *Tbr1^+/–^* mice. The VP is crucial for somatosensation and connections to somatosensory areas (SS) ^52^. Given that numbers of Thy1-YFP neurons were increased in both the primary and supplemental somatosensory cortices (SSp and SSs, respectively) of *Tbr1^+/–^* mice, we postulated that *Tbr1* deficiency prompts more innervation from SS to the VP. To explore this possibility, we performed retrograde labeling on the VP of *Tbr1^+/–^* mice using cholera toxin subunit B conjugated with Alexa Fluor 488 (CTB-488) (**Figure 5A**). After confirming the VP injection sites (**Supplemental Figure 5A-5C**), registration, segmentation and quantification were performed to analyze the distribution patterns of CTB-488^+^ cells (**Figure 5B-5D**). For comparison among animals, retrograde signals were normalized against the signal of the injection site (**Supplemental Figure 5D**). We identified several regions containing retrograde CTB-488 signals (**Figure 5C, 5D**). Among them, the ipsilateral SSp-ul (primary somatosensory area, upper limb), SSs and somatomotor network (MO) exhibited statistically significant differences between WT and *Tbr1^+/–^* mice (**Figure 5D, 5E, Supplemental Table 4**). Notably, the ipsilateral SSp-ul presented the greatest difference, both in terms of absolute CTB-488^+^ cell number and statistical significance, among the regions we considered (**Figure 5D, 5E, Supplemental Table 4**). Thus, there are more connections between the VP and somatosensory cortex in *Tbr1^+/–^* mice, supporting that *Tbr1* haploinsufficiency causes hyperconnectivity between the VP and SS.

**Figure 5.**
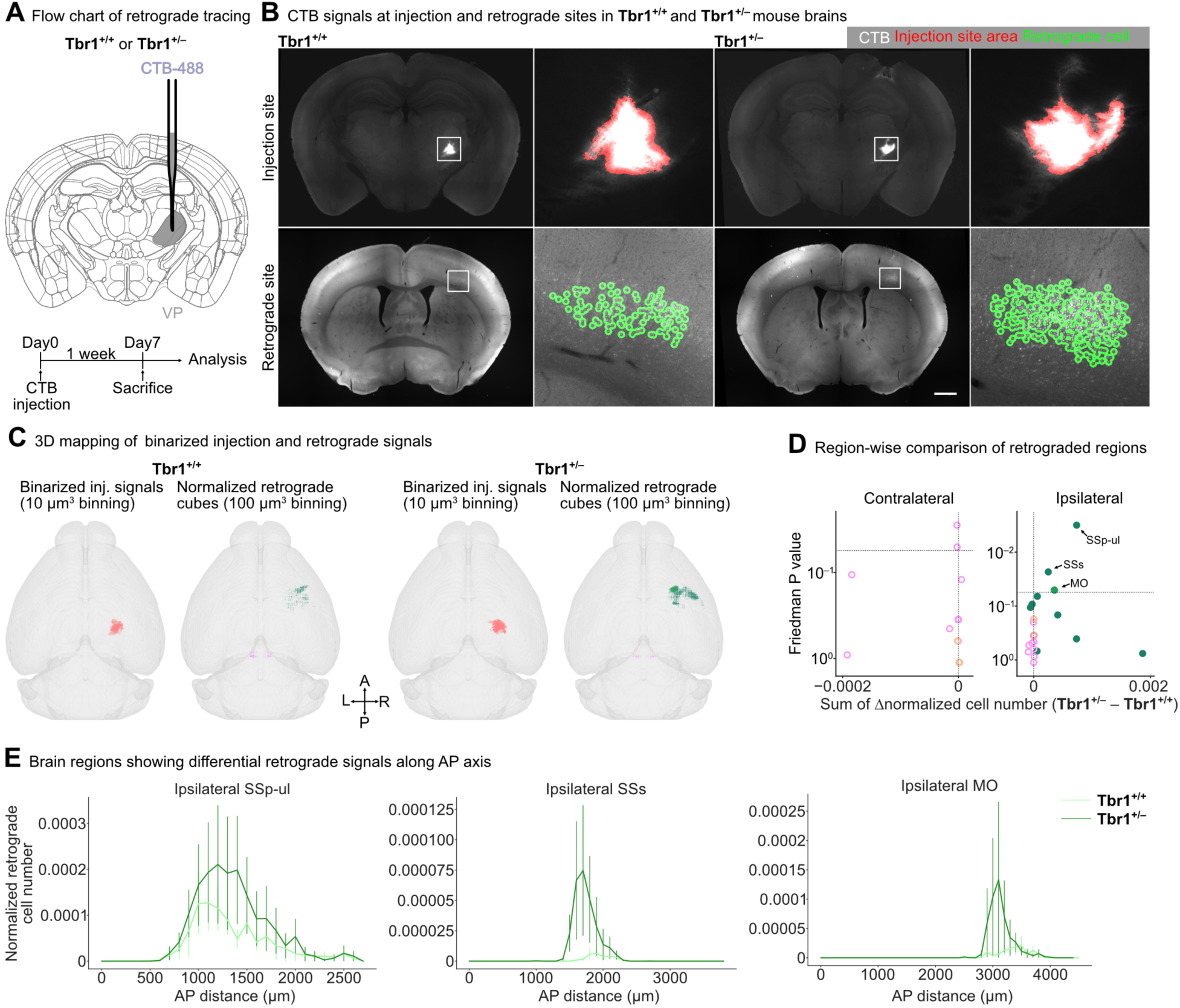
Hyperconnectivity to the VP in *Tbr1^+/^*^-^ mice. (**A**) Schematic of the experimental design (*Tbr1^+/+^*: n = 7; *Tbr1^+/–^*: n = 7). Brain samples injected with CTB-488 at the VP region were collected to trace signals in upstream brain regions. (**B**) Representative examples of retrograde tracing in *Tbr1^+/+^* and *Tbr1^+/-^* mice. Top, the VP injection site signals (white) were extracted via pixel classification, followed by object classification (red). Bottom, the retrograde signals (white) at the cerebral cortex were identified by the spot detection function of Imaris (green). Scale bar, 1 mm. (**C**) 3D visualization of binarized injection (red) and normalized retrograde (green) signals. A, anterior; P, posterior; L, left; R, right. (**D**) Structure-wise comparison of retrograde cell numbers between the *Tbr1^+/+^* and *Tbr1^+/–^* mice. Friedman’s one-way repeated measures were used for statistical analysis. The data points represent the brain regions. Closed circles, brain regions with manual-ROI correction; open circles, brain regions with auto-ROI correction. (**E**) The distribution of normalized retrograde signals along the anterior-posterior (AP) axis in the ipsilateral SSp-ul, SSs and MO regions. Data is represented as mean ± SEM. VP, ventral posteromedial nucleus; SSp-ul, primary somatosensory area, upper limb; SSs, supplemental somatosensory area; MO, somatomotor areas.

### Differential Thy1-YFP innervations

To systematically analyze the influence of *Tbr1*, *Nf1* and *Vcp* deficiencies on axonal projections of Thy1-YFP neurons, we screened the differentially innervated regions of Thy1-YFP neurons based on the correlation between the YFP^+^ cell numbers and total YFP signals in all brain regions (**Supplemental Figure 6**). We assumed that, for a given region, if the enhanced YFP signals are attributable to the increased Thy1-YFP cell population, we would detect a significant correlation between Thy1-YFP cell numbers and total YFP signals. Accordingly, significantly correlated regions would be less likely to display noticeably differential axonal innervation from upstream regions. Nevertheless, even if those regions were differentially innervated, YFP^+^ axons may be in the minority and therefore would be difficult to validate. Consequently, we aimed to identify noticeably differentially-innervated regions presenting no significant correlation between Thy1-YFP cell numbers and total YFP signals. After systematic screening (**Supplemental Figure 6**), we uncovered that regions meeting that criterion mainly lay in the thalamus (TH), hypothalamus (HY), midbrain (MB), and hindbrain (HB) of *Tbr1^+/–^*, *Nf1^+/–^* and *Vcp^+/R95G^* mice (**Figure 6, Supplemental Figure 7**). We also identified some regions in the cerebral cortex and cerebral nuclei, albeit not many (**Figure 6, Supplemental Figure 7**). These differentially-innervated regions are labeled with green font in **Figure 6**.

**Figure 6.**
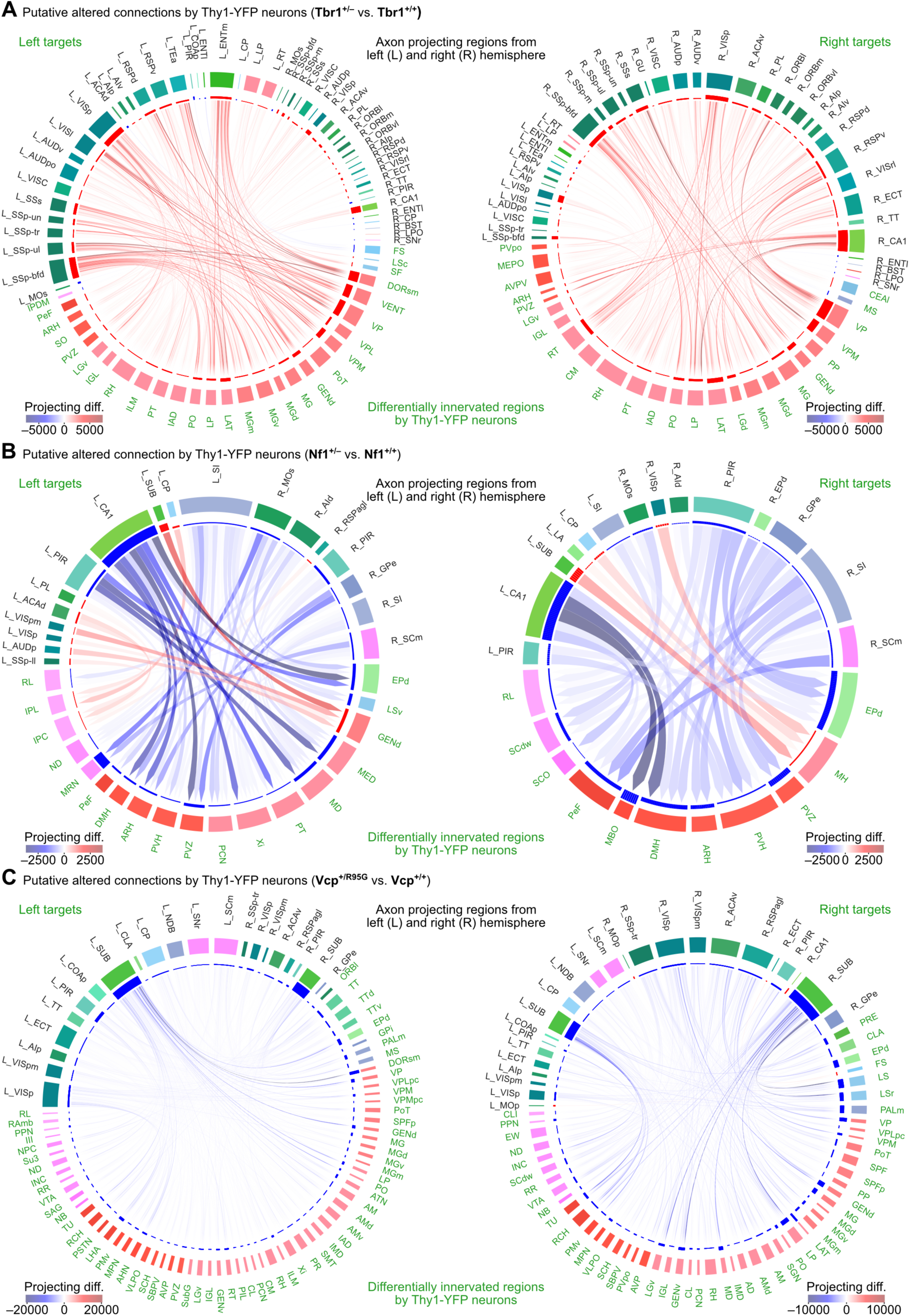
Prediction of differential Thy1-YFP circuits. (**A**) *Tbr1*^+/–^ vs. *Tbr1*^+/+^. (**B**) *Nf1*^+/–^ vs. *Nf1*^+/+^. (**C**) *Vcp*^+/R95G^ vs. *Vcp*^+/+^. The differential Thy1-YFP circuits of the above three groups were predicted based on total YFP signals, Thy1-YFP cell numbers, and the Allen Mouse Brain Connectivity Atlas. The differentially-innervated target regions in the left and right hemispheres are indicated with green font and are listed outside the circos plots in the left and right panels, respectively. Names of the axon-projecting regions in the left (L_) or right (R_) hemisphere (black font) are also indicated. The segments of the outer circles represent the color codes of brain regions, as detailed in Figs. 3 and 4. The segments of the inner circles represent the normalized sum of ΔYFP signals at innervated regions or the normalized sum of ΔYFP^+^ cell numbers at axon-projecting regions. The lines within the circus plots connect the axon-projecting regions (blunt end) to the innervated regions (pointed end). Heatmaps of line colors in the bottom right or left corners of each plot represent the projecting differences having differential tendencies (red: mutant > WT, blue: mutant < WT). The thickness of the lines corresponds to the absolute value of the log-transformed projecting difference.

In *Tbr1^+/–^* mice, these differentially-innervated regions exhibited more extensive Thy1-YFP innervation (**Figure 6, Supplemental Figure 7**), which was not the case for *Nf1^+/–^* or *Vcp^+/R95G^* mice (**Figure 6, Supplemental Figure 7**). Among these regions, we noticed that only the GENd (Geniculate group, dorsal thalamus) was shared among all three groups (**Supplemental Figure 7D, 7E, 7F,** underlined), albeit with distinct differences. For instance, the GENd of the left and right hemispheres of *Tbr1^+/–^* mice had greater Thy1-YFP innervation, yet fewer innervations in both hemispheres of the GENd in *Vcp^+/R95G^* mice. Moreover, only the GENd of the left hemisphere of *Nf1^+/–^* mice exhibited increased Thy1-YFP innervation (**Supplemental Figure 7D, 7E, 7F,** underlined). Since the GENd is known to form reciprocal connections with visual areas (VIS), the altered Thy1-YFP innervation at the GENd indicates changes in VIS. Consistent with this speculation, we observed that both *Tbr1^+/–^* and *Nf1^+/–^* mice possessed more Thy1-YFP neurons at the VISp than their respective WT littermates (**Figure 4A-4D**), whereas *Vcp^+/R95G^* mice had fewer Thy1-YFP neurons at the VISp (**Figure 4E-4F**). These results reveal altered innervations by Thy1-YFP neurons under different mutation scenarios.

### Predicting altered Thy1-YFP circuits

Next, we examined the potential upstream regions of all differentially Thy1-YFP-innervated regions. To do so, we used the Allen Mouse Brain Connectivity Atlas and our datasets of YFP signals and YFP^+^ cell numbers (**Figure 4, Supplemental Figure 7 and Tables 5-7**) to predict potential alterations of neural connections in *Tbr1^+/–^*, *Nf1^+/–^* and *Vcp^+/R95G^* mice (see **Supplemental Figure 8** and **Methods** for details). Our results show that *Tbr1* deficiency generally enhanced innervations by Thy1-YFP projection neurons into the TH, HY, MB and HB regions (**Figure 6A**), including hyperconnectivity between the SS and VP, which was already validated by retrograde labeling (**Figure 5**). *Nf1* deficiency had a mixed effect on Thy1-YFP neuron innervation, with some connections predicted as being enhanced but a majority was reduced (**Figure 6B**). Thy1-YFP neuronal innervations were generally reduced for *Vcp^+/R95G^* mice (**Figure 6C**). These results reveal differential impacts of the *Tbr1*, *Nf1* and *Vcp* mutations on axonal innervation by Thy1-YFP neurons.

### Targeting of the piriform complex and other sensory regions

The above-described results reveal three characteristics of the Thy1-YFP connections in *Tbr1^+/–^*, *Nf1^+/–^* and *Vcp^+/R95G^* mice. First, general patterns of differential YFP signals and YFP^+^ cell numbers varied among these ASD mouse models (**Figure 3, 4**), resulting in differential alterations of neural connectivity, at least for the Thy1-YFP neurons in the three mutant mouse lines **(Figure 6, Supplemental Figure 7**), and reflecting the divergent effects of various autism-linked mutations on Thy1 projection neurons. Second, we noticed that ΔYFP signals and ΔYFP^+^ cell numbers frequently presented left-right asymmetries for all three mouse lines (**Figure 3, 4, 6, Supplemental Tables 5-7**). Among all six group comparisons shown in **Figures 3** and **4**, we found that the CA1 was the only region showing left-right asymmetry in terms of YFP^+^ cell numbers, albeit with slight differences. *Tbr1^+/–^* and *Vcp^+/R95G^* mice both tended to have more Thy1-YFP neurons in the right CA1 (**Figure 4A, 4B, 4E, 4F**), whereas the left CA1 of *Nf1^+/–^* mice had fewer Thy1-YFP neurons (**Figure 4C, 4D**). However, based on the predicted neuronal circuits (**Figure 6**), this left-right asymmetry may not greatly impact CA1 connectivity in *Vcp^+/R95G^* mice, yet it is overwhelmingly dominant in *Nf1^+/–^* mice (**Figure 6B, 6C**). Thus, the patterns of left-right asymmetry also varied among the three mouse lines. Finally, among the more than 300 brain regions we examined, only one (i.e., the piriform area; PIR), exhibited the same change in YFP signal and YFP^+^ cell number in all three types of mutant mice (**Figure 3, 4, Figure 7A**, 7B). Based on gene expression data from the Allen Mouse Brain Atlas (https://mouse.brain-map.org), *Tbr1*, *Nf1* and *Vcp* are all expressed in the PIR (**Supplemental Figure 9A**). We observed that Thy1-YFP neurons are mainly present in the deep layer (i.e., layer III) of the PIR (**Supplemental Figure 9B**). Notably, numbers of Thy1-YFP projection neurons and total YFP signals were reduced in the PIR of both the left and right brain hemispheres (**Figure 3, 4, 7A, 7B, Supplemental Figure 10-12**), implying potentially impaired PIR function in our mutant mice.

**Figure 7.**
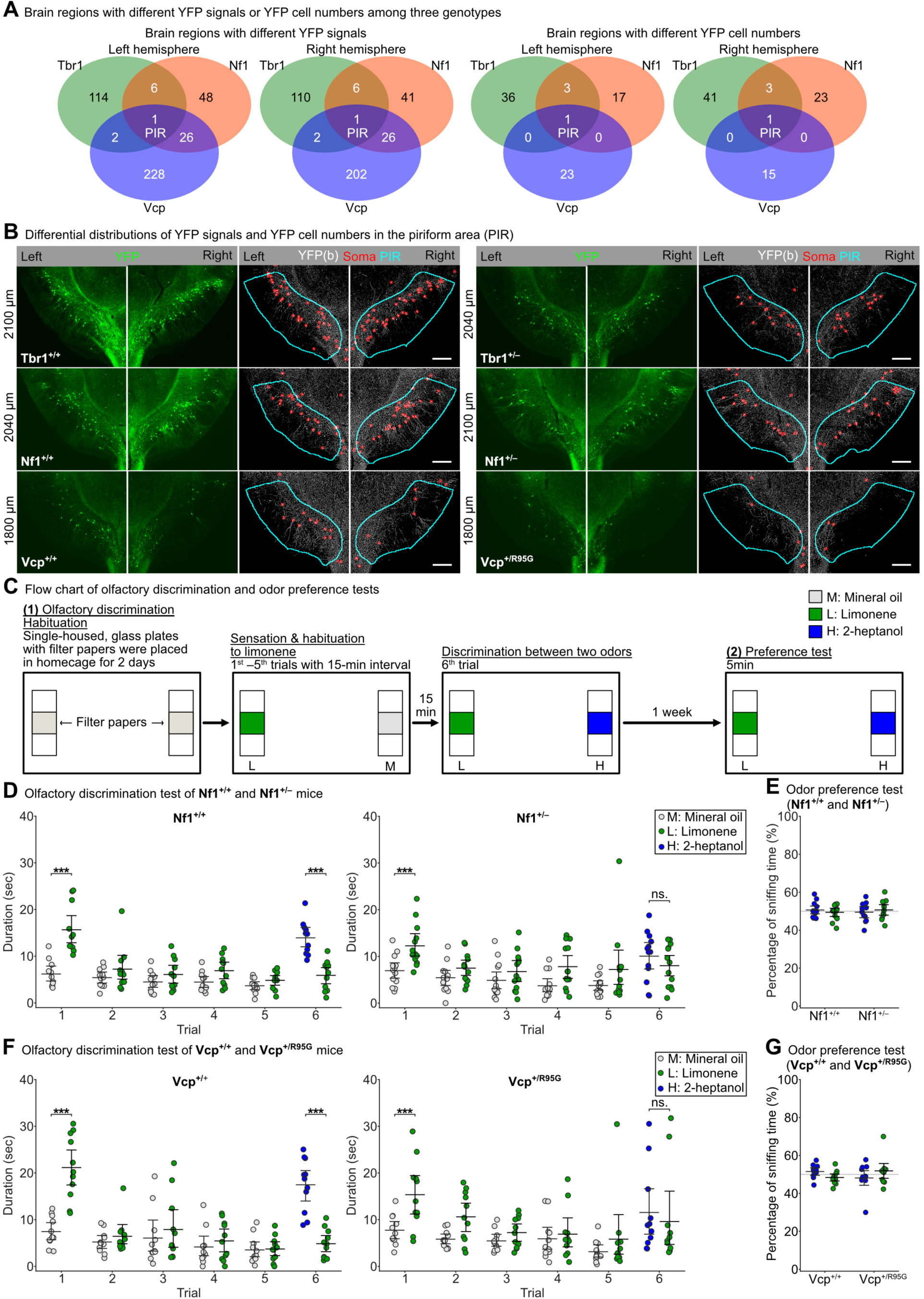
Impaired olfactory discrimination in *Nf1^+/–^* and *Vcp^+/R95G^* mice. (**A**) Venn diagrams showing the number of brain regions with differential YFP signals (left panels) and YFP cell numbers (right panels) specific to each mutant genotype or common among two or three genotypes (*Tbr1*^+/–^ vs. *Tbr1*^+/+^, *Nf1*^+/–^ vs. *Nf1*^+/+^, *Vcp*^+/R95G^ vs. *Vcp*^+/+^). Only the PIR region shows a common deficiency in YFP signals and YFP^+^ cell numbers among the three mutant genotypes. (**B**) Registered Thy1-YFP images (green signals) and their binarized YFP images (white) with detected YFP^+^ soma (red) indicating that the mutant mouse brains of three genotypes (right example) have fewer YFP^+^ cells in the PIR (outlined by cyan frame) relative to their WT littermates (left example). (**C**) Flow chart of behavioral assays, including the olfactory discrimination and olfactory preference tests. A one-week interval separated the olfactory discrimination and odor preference tests. Limonene and 2-heptanol represented test odors, and mineral oil acted as a control. (**D**) Results of the olfactory discrimination for *Nf1^+/+^* and *Nf1^+/–^* mice. (**E**) Results of the odor preference tests for *Nf1^+/+^* and *Nf1^+/–^* mice. (**F**), (**G**) Results of the olfactory discrimination and odor preference tests for *Vcp^+/+^* and *Vcp^+/R95G^* mice. Data is represented as mean ± SEM with individual sample points in (**D-G**). Sample sizes: *Nf1^+/+^* n = 11; *Nf1^+/–^* n = 13; *Vcp^+/+^* n = 11; *Vcp^+/R95G^* n = 11. Scale bar in (**B**), 300 µm.

In addition to the PIR, we found that some sensory-related brain regions also exhibited changes in the mutant mice. For instance, both total YFP signal and YFP^+^ cell number of the VISp were increased in *Tbr1^+/–^* and *Nf1^+/–^* mice, but they were reduced in the *Vcp^+/R95G^* mice (**Figure 3, 4**). In terms of the MO, total YFP signals were upregulated in *Tbr1^+/–^* mice, yet downregulated in both *Nf1^+/–^* (only in the left hemisphere) and *Vcp^+/R95G^* mice (**Figure 3**). Moreover, both *Tbr1^+/–^* and *Nf1^+/–^* mice had more YFP^+^ cells in the SSp, which was not the case for *Vcp^+/R95G^* mice (**Figure 4**). Thus, our analyses indicate that cortical regions controlling olfaction, vision, the somatomotor network, somatosensation, and the thalamic region related to sensory gating are particularly susceptible to structural changes in diverse ASD conditions, with the PIR being the only region in this respect shared by all three mutant mouse lines we considered.

### Impairment of olfactory discrimination

The PIR is the olfactory cortex involved in high-order processing of olfactory information ^53,54^. Our previous study showed that *Tbr1* haploinsufficiency impairs neuronal activation of the olfactory system, including the PIR, in response to olfactory stimulation, resulting in defective olfactory discrimination in mice ^28^. That finding echoes the deficits of Thy1-YFP projection neurons in the PIR revealed in our current report.

Given that *Nf1^+/–^* and *Vcp^+/R95G^* mice shared the same PIR phenotype as *Tbr1^+/–^* mice, we speculated that *Nf1^+/–^* and *Vcp^+/R95G^* mice also likely exhibit an impaired olfactory response. In our previous study ^28^, we used two distinct odors, i.e., limonene and 2-heptanol, to test the olfactory responses of *Tbr1^+/–^* mice. In that study, *Tbr1^+/–^* mice did not present any preference for limonene or 2-heptanol, i.e., equivalent to WT littermates. However, unlike WT littermates, *Tbr1^+/–^* mice could not discriminate between limonene and 2-heptanol ^28^. We subjected *Nf1^+/–^* and *Vcp^+/R95G^* mice sequentially to the same olfactory analyses, i.e., olfactory discrimination and odor preference tests (**Figure 7C**). In the first trial of the olfactory discrimination test, *Nf1^+/–^* mice and their WT littermates responded normally to limonene (**Figure 7D**). They also presented a desensitized response for subsequent trials 2 to 5 (**Figure 7D**). However, upon encountering both limonene and the novel odor 2-heptanol in trial 6, only WT littermates presented a preference for 2-heptanol, with *Nf1^+/–^* mice incapable of recognizing 2-heptanol as a new odor and consequently not showing a preference for it in trial 6 (**Figure 7D**). Both *Nf1^+/–^* mice and WT littermates exhibited a comparable tendency to sniff limonene and 2-heptanol in the preference test (**Figure 7E**). Thus, our *Nf1^+/–^* mice exhibited impaired olfactory discrimination and this deficit was not caused by a different odor preference.

In terms of *Vcp^+/R95G^* mice, although individual variation in our olfactory analyses appeared higher than for the other two mutant lines, our results also revealed that the *Vcp* mutation impaired olfactory discrimination but did not influence odor preference (**Figure 7F, 7G**). Thus, taken together with the results of our previous study ^28^, all *Tbr1^+/–^*, *Nf1^+/–^* and *Vcp^+/R95G^* mice exhibit impaired olfactory discrimination. In fact, impaired olfactory processing and a defective piriform cortex are common deficiencies shared among *Tbr1^+/–^*, *Nf1^+/–^* and *Vcp^+/R95G^* mice.

## Discussion

We have used a mesoscale system to analyze Thy1-YFP reporter patterns in three ASD mouse models. By deploying the BM-auto platform to enhance accuracy and efficiency, our quantification analysis reveals distinct innervation patterns of Thy1-YFP projection neurons and differential alterations of Thy1-YFP neuronal density in the *Tbr1^+/–^*, *Nf1^+/–^* and *Vcp^+/R95G^* mice. Despite the diverse outcomes, importantly we noted that the piriform cortex exhibits reductions in both YFP signal and Thy1-YFP cell number in all three of the mutant mouse lines, in both the left and right brain hemispheres. Moreover, all three of our ASD models displayed defective olfactory discrimination. In addition to the piriform cortex, we identified other sensory areas, such as the VIS, SS, and GENd, which exhibited alterations in Thy1-YFP patterns, though the tendencies differed or changes were uncovered in two but not all three mutant lines. Nevertheless, our results strongly indicate that sensory impairment, especially an abnormal olfactory response, is common under different ASD conditions, supporting the sensory-first hypothesis of developing ASD ^55–58^. ASD is defined by a dyad of impairments: (1) defective social interaction and communication; and (2) repetitive and restricted interests and abnormal sensations ^59,60^. Given that defective social interaction and communication are explicit behavioral deficits, social deficits have garnered much attention in studies of ASD. However, since sensory processing is crucial for developing cognitive systems, alterations in sensory processing have been proposed to contribute to ASD ^55–57^. Certain studies of some ASD models, including *Cntnap2, En2, Fmr1, Gabrb3*, *Mecp2*, *Shank2*, *Shank3*, and *Syngap1* mutant mice, have shown defective tactile and pain sensations as they develop autism-like behaviors ^61–67^. In our current study, we also found that the SSp exhibits divergent YFP signals in all three mutant mouse lines we considered, being upregulated in both *Tbr1^+/–^* and *Nf1^+/–^* mice, yet downregulated in *Vcp^+/R95G^* mice. This outcome implies altered mechanosensation in the three mutant lines, though further studies are necessary to provide direct evidence.

Importantly, we have shown that the olfactory piriform cortex exhibits identical cellular and functional deficits in all three ASD models. Olfactory sensation is developed and becomes functional during the embryonic stage ^68,69^. Fetuses receive and process olfactory information in the womb and even develop a preference for the respective odors after birth ^69–71^. The piriform cortex is embryonically active, and this early activation is required for cortical connectivity during development ^72^. In addition, olfactory responses at the developmental stage appear to be involved in building up multiple brain networks to detect environmental information, influencing eating behaviors and nutrient intake, and regulating social communication ^70,73^. In a cohort study of ASD patients, typically developing controls reduced their sniff response to unpleasant odors, but the subjects suffering ASD exhibited similar sniff responses to both pleasant and unpleasant odors ^74^. This finding is very similar to the impaired olfactory discrimination we detected in the three ASD models we tested in the current study, confirming the idea that impaired olfaction is a common feature in ASD.

Combining our datasets and information from the Allen Mouse Brain Connectivity Atlas, we predicted altered innervation patterns by Thy1-YFP neurons in the brain, with *Tbr1*, *Nf1* and *Vcp* deficiencies having distinct impacts on axonal projection. We also found that these three ASD-associated genes have different effects on the numbers of Thy1-YFP neurons, indicating that ASD-linked genes control the complex processes of neuronal development or maintenance. Notably, these effects appear to be non-cell-autonomous since Tbr1 is highly expressed in layer 6 cortical neurons rather than layer 5 Thy1^+^ neurons ^34^. Nevertheless, our whole-brain analysis proved sufficiently powerful to reveal differences.

We adopted the Thy1-YFP-H line as a reporter to analyze the effects of *Tbr1*, *Nf1* and *Vcp* deficiencies on neural connectivity. In the cerebral cortex, the Thy1-YFP transgene is primarily expressed in layer 5 projection neurons, which project to ipsilateral subcortical regions, so it is suitable for analyzing connections between the cortical and subcortical regions. To examine other types of connections, such as contralateral projections between two brain hemispheres, other reporter mice will be needed. Even for layer 5 cortical neurons, the Thy1-YFP transgene is only expressed in a subpopulation, i.e., not in all of them. Thus, it would also be interesting to investigate the projection patterns of those Thy1-YFP negative neurons in the future. Apart from being applied to ASD mouse models, our BM-auto platform can also be employed in other studies, greatly accelerating whole-brain analyses for various research goals.

## Resource availability

### Lead contact

Requests for further information and for resources and reagents should be directed to and will be fulfilled by the lead contact, Dr. Yi-Ping Hsueh

### Material availability

This study did not generate new unique reagents.

### Data and code availability

All Supplemental Videos and Tables have been deposited at Zenodo (https://zenodo.org) and are publicly available as of the date of publication at (DOI). Custom codes used in the current study have been uploaded to GitHub (https://github.com/genephil/dlreg and https://github.com/HsuehYiPing/Thy1YFP and) and Zenodo (DOI:)

## Acknowledgments

We thank Prof. Mark Liao at the Institute of Information Science, Academia Sinica, and Profs. Anne Calof and Arthur Landers at UC, Irvine, for helpful discussion and suggestions; the Allen Institute (http://portal.brain-map.org) for its open resource mouse brain atlas; the Imaging Core and Animal Facility of the Institute of Molecular Biology, Academia Sinica, for technical assistance; Dr. John O’Brien for English editing; and members of Y.-P.H.’s laboratory for technical assistance and discussion. This work was supported by grants from Academia Sinica (https://www.sinica.edu.tw, AS-IA-111-L01 and AS-TP-110-L10 to Y.-P.H.), and the National Science and Technology Council (https://www.nstc.gov.tw/, NSTC 112-2326-B-001-008 to Y.- P.H., NSTC 109-2628-M-001-001-MT4 to B.-C.C.). The funders had no role in study design, data collection and analysis, the decision to publish or the preparation of the manuscript.

## Author contributions

Conceptualization, T.-T.H. and Y.-P.H.; Methodology and investigation, T.-T.H., M.-H.L., T.- N.H., Chung-Yu Wang, C.-M.-L. and B.-C.C.; Software, T.-T.H., C.-P.C., M.-H.L. and Chien-Yao Wang; Writing – original draft, T.-T.H., M.-H.L., T.-N.H., C.-P.C., Chien-Yao Wang, B.-C.C. and Y.-P.H.; Writing – review and editing, T.-T.H., M.-H.L., T.-N.H., C.-P.C., Chien-Yao Wang, B.-C.C. and Y.-P.H.; Funding acquisition, Y.-P.H.; Supervision and project administration, Y.-P.H. All authors contributed to the article and approved the submitted version.

## Supplemental information

Supplemental Figures can be found online. All Supplemental Videos 1-6 and Supplemental Tables 1-8 are available at Zenodo (DOI:).

## Materials and Methods

### Ethics statement

All animal experiments were performed with the approval of the Academia Sinica Institutional Animal Care and Utilization Committee and in strict accordance with its guidelines (Protocol No. 18-10-1234 and 23-03-1990). Human subjects were not included in the current work.

### Mice

The *Tbr1*^+/−^ mice were originally provided by Dr. J. L. Rubenstein (Department of Psychiatry, University of California, San Francisco) ^75^. The *Nf1*^+/−^ mice (B6.129S6-Nf1^tm1Fcr^/J, Strain No: 002646) ^76^ and Thy1-YFP-H transgenic mice (B6.Cg-Tg(Thy1-YFP)HJrs/J, Strain No: 003782) ^31^ were purchased from Jackson Laboratory. We use “Thy1-YFP” mice to denote Thy1-YFP-H mice in the current report. The *Vcp*^+/R95G^ knockin mice were generated and characterized previously ^30,40^. All mouse lines were maintained by backcrossing to WT C57BL/6 mice. To introduce the Thy1-YFP transgene into ASD model mice, *Tbr1*^+/–^, *Nf1*^+/–^ and *Vcp*^+/R95G^ mice were crossed with the Thy1-YFP-H (Thy1-YFP) mice. Male *Tbr1*^+/–^;*Thy1-YFP^+/–^*, *Nf1*^+/–^;*Thy1-YFP^+/–^* and *Vcp*^+/R95G^;*Thy1-YFP^+/–^* mice were subjected to whole-brain imaging because previous studies have shown ASD-linked deficits in male mutant mice ^28–30,77^. Given that the backcross generations differed among these three mouse lines, male WT littermates of each line, i.e., *Tbr1*^+/+^;*Thy1-YFP^+/–^*, *Nf1*^+/+^;*Thy1-YFP^+/–^* and *Vcp*^+/+^;*Thy1-YFP^+/–^*, were used for comparison with the corresponding mutant mice. All animals were housed in the animal facility of the Institute of Molecular Biology, Academia Sinica, under controlled temperature and humidity and a 12 h light/12 h dark cycle. The experiments were performed at 3-4 months of mouse age.

### Brain sample preparation and light-sheet microscopy imaging

*Tbr1^+/+^* and *Tbr1^+/–^* mice containing a single copy of the Thy1-YFP transgene were subjected to analyses. Mice were deeply anesthetized via intraperitoneal (i.p.) administration of 2,2,2-Tribromoethanol (0.7-0.8 mL per mouse), and they were sacrificed by perfusing them sequentially with phosphate-buffered saline (PBS) and 4% paraformaldehyde (PFA) in PBS by means of a peristaltic pump (101U/R, Watson-Marlow, flow rate: 1.5 mL/min). The brains were dissected out and continuously immersed in 4% PFA for post-fixation at 4 °C for 72 hours with gentle horizontal shaking (10 rpm). Brain samples were then washed with PBS three times, before being subjected to tissue clearing. We used the PEGASOS tissue clearing method ^47,78^ involving a passive immersion procedure to prepare brain samples for light-sheet imaging as follows. First, the brain samples were sequentially decolorized with 25% w/v Quadrol (Sigma-Aldrich, 122262) in H2O for 2 days and then with 3% ammonium solution (freshly prepared, v/v with H2O) for 1 day. The decolorizing procedures were conducted at 37 °C with shaking (10 rpm rocking shake). Second, the samples were washed with PBS three times, before undergoing gradual delipidation by sequentially incubating them in a 30%, 50%, and 70% gradient of tert-Butanol (tB, Sigma-Aldrich, 360538) delipidation solutions (various % v/v tB, 3% w/v Quadrol in H2O) for 8 hours, 1 day and 2 days, respectively. Gradual delipidation was conducted at 37 °C with shaking (10 rpm rocking shake). Third, after delipidation, the samples were dehydrated using tB-PEG dehydration solution (27% v/v PEG methacrylate Mn 500 (Sigma-Aldrich, 447943), 70% v/v tB, 3% w/v Quadrol) for 2 days at 37 °C with shaking (10 rpm rocking shake). Fourth, the dehydrated brains were cleared using BB-PEG clearing solution (75% v/v benzyl benzoate (Sigma-Aldrich, B6630), 22% v/v PEG methacrylate Mn 500, 3% w/v Quadrol) for 2 days at 37 °C with 10 rpm rocking shaking. Finally, the cleared brain samples were immersed in another 100 mL tB-PEG clearing solution for refractive index (RI) matching. RI matching was continued for at least 1 week at room temperature before light-sheet imaging. The BB-PEG clearing solution used for RI matching was also used for light-sheet imaging. The cleared mouse brain was translated to multiple positions using a “snake-by-row right-then-down” strategy to image the whole brain sequentially in 3D. The dual-side Bessel illumination light from two excitation objectives and the fluorescence signals were collected by an Olympus 4X NA 0.28 dry objective and projected onto a sCMOS Camera, Andor Zyla4.2 plus. We utilized Fiji’s Grid/Collection stitching feature to stitch the data into a complete digital image of the mouse brain.

### Mesoscopic BM-auto pipeline

To compare the YFP-labeled circuits between mutant and WT mice, we conducted a mesoscopic whole-brain scale analysis as described previously ^24^ with some modifications (**Figure 2, Supplemental Figure 1-3**). Details of the procedures on the 3-4-month-old male mice are described in the following sections.

#### Step1: Brain sample preparation and high-content immunofluorescence imaging

Mice were deeply anesthetized via i.p. administration of 2,2,2-Tribromoethanol (0.7-0.8 mL per mouse), and they were sacrificed by perfusing them with PBS, followed by 4% PFA in PBS. The brains were dissected out and continuously immersed in 4% PFA for post-fixation overnight (∼16 hours) at 4 °C. After post-fixation, brains were dehydrated in 30% sucrose in PBS at 4 °C for 2 days and then embedded in OCT (4583, Tissue-TeK). We cryosectioned each brain from the posterior to anterior cerebrum into at least 120 consecutive coronal brain slices of 60 µm thickness using a cryostat microtome (CM1900, Leica). The cerebellum and olfactory bulb were excluded from our analysis. The coronal brain sections were washed with Tris-based saline (TBS, 25 mM Tris-Cl pH 7.5, 0.85% NaCl) three times to remove residual OCT and stained with DAPI (1 µg/mL, ThermoFisher Scientific). The processed brain sections were imaged with a high-content imaging system (Molecular Devices) hosting a 4x objective. Thy1-YFP signals, DAPI signals, and autofluorescence tissue background were acquired with different fluorescence filter sets. Images from one brain were stitched together using MetaXpress 6.2.3.733 (Molecular Devices) and manually aligned in Amira 6.4 (ThermoFisher Scientific) or Etomo ^79^ according to DAPI signals.

#### Step2: Registration to Allen Mouse Common Coordinate Framework version 3 (CCFv3) and auto-*ROI correction*

(1) Initial Registration (Reg)

After acquiring the raw images of aligned consecutive slices, we registered the image series to CCFv3 ^80^. We downloaded high-resolution coronal templates with a 10 μm resolution along the anterior-posterior (AP) axis using Allen Software Development Kit (Allen SDK) Python code and then reduced the image size to fit the high-content images (6480 x 4547 pixels). According to the autofluorescence tissue background of the raw images, we manually matched the corresponding template to each raw image according to specific landmark brain structures, such as the beginning and end of the hippocampus (HIP), the location of the anterior commissure (act and aco), and the disconnections of the corpus callosum (cc). After determining the matched template for each raw image, binary experimental images were registered to the matched binary templates utilizing the medical image registration library SimpleElastix ^81^, first with 12 global (translation and affine) and then one local (B-spline) transforming parameters. Finally, the stored multiple transforming parameters were applied to the experimental images of Thy1-YFP and tissue background signals to generate registered images.

(2) Manual ROI corrections of 37 brain regional masks

We accessed the Reference Space of CCFv3 to obtain the coronal indicator masks of each brain region by using Allen SDK Python code (https://allensdk.readthedocs.io/en/latest/_static/examples/nb/reference_space.html). The CCFv3 regional masks could separately compartmentalize YFP pixels and YFP^+^ cells into different brain regions, which served as regions of interest (ROI) to quantify the binarized YFP pixel number (YFP signal) or YFP^+^ cell number for each brain region along the AP axis. Before quantification, we performed ROI correction on 37 selected brain regions in both hemispheres of each brain sample, as described previously ^24^. In brief, each brain region’s original coronal indicator masks were separated into left and right hemispheric masks and converted to ImageJ ROIs by sequentially applying the ImageJ Create Selection function and adding it to ImageJ ROI Manager. Using the initially registered images with tissue background or Thy1-YFP signals as references, ROIs were verified and corrected by manually adjusting their shapes and locations with ImageJ tools. After ROI corrections, the ROIs of each brain region in both hemispheres were converted back to image masks by applying the ImageJ Create Mask function. The manually corrected indicator masks were then used to develop a deep learning model for auto-ROI correction of CCFv3 regional masks.

(3) Deep learning model for CCFv3-to-image registration (DeepReg)

We developed a deep learning model called DeepReg, aimed at enhancing whole-brain registration by leveraging annotations from limited brain regions (37 selected brain regions). DeepReg takes the CCFv3 template and our brain image as inputs to estimate a dense, spatial deformation field from CCFv3 for the images. To address the challenge of modality differences between CCFv3 and our brain images, we introduced a dual-stream U-Net architecture, pre-trained respectively on the annotated brain regions, to project both the template and images into a shared semantic space for modality alignment. The image features extracted by this pre-trained dual-stream encoder were then passed through a correlation-aware coarse-to-fine registration decoder ^82^, producing a deformation field represented by band-limited field representations ^83^. Through a weakly supervised learning approach, the model achieves coarse-to-fine high-resolution smooth deformation and accurate semantic segmentation of the annotated brain regions.

For our experiments, we randomly divided our data into training (30 images), validation (4 images), and testing sets (4 images). During model training, we applied weakly supervised learning by utilizing images and their manually corrected indicator masks from the training set. Unlike unsupervised learning, this approach features an additional soft Dice loss term, which improves the model’s ability to learn and accurately identify anatomical structures in the brain images. After training, we evaluated the performance of our deep learning model on the validation set, focusing on the 37 selected brain regions, as well as other brain regions.

(4) Auto-ROI correction of original CCFv3 regional masks by inverse refinement

First, deformation fields for auto-ROI correction were obtained by DeepReg model-based reverse registration of original CCFv3 templates (1140 x 800 pixels) to initially registered images with tissue background. The corresponding deformation fields were used to inversely refine the original CCFv3 regional masks to auto-corrected CCFv3 regional masks, which were then used for quantification.

#### Step3: Image segmentation for Thy1-YFP signals and detection of YFP-positive soma with strong signals (also see **Supplemental Figure 3**)

Before quantifying and comparing Thy1-YFP signals, we needed to eliminate the variance of YFP fluorescence signals caused by experimental variation or tissue background interference. Image segmentation of YFP signals that determines which pixels are YFP-positive was performed as described previously (**Supplemental Figure 3,** lower panel) ^24^. First, we deployed the pixel classification algorithm ilastik1.3.3post3 ^84^ to predict the probability that every pixel in a registered image displayed YFP signals. The YFP probability maps of each image were cross-referenced against the registered images pixel by pixel. The resulting filtered images (raw signal intensity x probability) were further processed to intensify edges by subtracting the Gaussian blurred version of the filtered images. The edge-enhanced filtered images were then subjected to thresholding/binarization (> 6x S.D. of the Z-stacked image with filtering and edge enhancement) to isolate YFP-positive pixels.

We further mapped the locations of soma displaying strong YFP signals in each image. The results indicated the numbers of Thy1-YFP neurons. The images with YFP signals were preprocessed by applying rolling-ball background subtraction (ImageJ, radius 50 pixels). The spot detection algorithm of Imaris 9.3 (Bitplane) was then applied to detect strong YFP^+^ cells from background-subtracted images using consistent criteria (Detection spot diameter: 15 µm; Detection threshold: 2 x S.D. of the Z-stacked image with background subtraction). The locations of YFP^+^ cells in each image could then be exported. The images with binarized YFP pixels and the information of YFP^+^ cell locations were subjected to quantification.

#### Step4: Application of automatic and manually ROI-corrected brain regional masks to quantify YFP pixels (signal) and YFP^+^ cell numbers

The auto-ROI-corrected and 37 manually ROI-corrected CCFv3 regional masks were used to compartmentalize YFP pixels and YFP^+^ cells into different brain regions, before quantification of YFP signal and YFP^+^ cell numbers along the AP axis of each brain region was carried out. All computations were done using the NumPy, Pandas, OpenCV, scikit-image and GeoPandas Python packages. The data compartmentalizing YFP signal and YFP^+^ cells into different brain regions were subsequently used for 3D visualization. Data on quantified binarized YFP signal and YFP^+^ cell numbers at each slice of every brain region were used for further analyses (see below).

### Slice-based quantification and comparisons between WT and mutant mice

Slice-based quantifications involved in calculating the binarized YFP signal and YFP^+^ cell numbers along the AP axis were conducted for each brain region. Then, we carried out 100-µm binning along the AP axis for region-wise comparisons. For the three compared groups (*Tbr1^+/–^* vs. *Tbr1^+/+^*, *Nf1^+/–^* vs. *Nf1^+/+^*, *Vcp^+/R95G^* vs. *Vcp^+/+^*), we selected the brain regions containing the same number of binned slices in all compared group samples for analysis. The statistical significance of differences in YFP signal or YFP^+^ cell distributions was tested by Friedman’s one-way repeated measures analysis (utilizing the Pingouin Python package), followed by P value adjustment to control for the false discovery rate (using the statistical function false_discovery_control of the Scipy Python package). Then, we conducted pair-wise subtractions of YFP signal or YFP^+^ cell numbers for the same AP coordinates between all possible pairs. The differences along the AP axis of all sample pairs were then averaged and summed into single metrics (sum of ΔYFP or sum of ΔYFP^+^ cell number) to represent the extent of the differences.

### 3D visualization

#### Summarized 3D stack of binarized YFP images

To visualize the averaged differences of YFP signals between WT and mutant mice in three compared groups (*Tbr1*^+/–^ vs. *Tbr1*^+/+^, *Nf1*^+/–^ vs. *Nf1*^+/+^, *Vcp*^+/R95G^ vs. *Vcp*^+/+^), the binarized YFP image series of individual brains were binned along the dorso-ventral and medio-lateral axes into 10 µm/pixel. Each binned pixel intensity represented the Thy1-YFP probability. The AP coordinates of the binned YFP image series were converted into the respective CCFv3 coordinates (10 µm/pixel), and all binned YFP image series of individual brains were then merged. If the same AP coordinate contained multiple binned YFP image series from different brain samples, we averaged the intensity of those binned YFP image series. If an AP coordinate contained only one binned YFP image, that binned image represented the Thy1-YFP signals for that particular AP coordinate. The 3D stack of binned and averaged YFP image series was adopted to illustrate Thy1-YFP signals at a whole-brain scale, 3D mapping of Thy1-YFP signal differences between WT and mutant mice, and prediction of differential Thy1-YFP circuits (see below). Image processing was carried out using the Numpy and scikit-image Python packages and ImageJ.

#### 3D mapping of YFP signal or YFP^+^ cell distributions

The 3D mapping of YFP signal or YFP^+^ cell numbers was conducted as described previously ^24^. The CCFv3 whole brain shell was obtained from the image stacks of CCFv3 “root” coronal indicator masks and converted to an isosurface mesh with smoothing using the vedo Python package. To visualize the YFP signal distributions in different brain regions, binned and averaged YFP image stacks within particular regions were employed, and binned pixels (10 µm^3^ cubes) were converted into vectorized points using Numpy, Pandas and vedo. Each vectorized point expressed the information of the brain region to which the point belonged (discontinuous color map) and YFP probability (alpha level). Differential YFP signals were generated by subtraction of the YFP probability of each vectorized point in the same location between the averaged mutant and WT brains. Scaled differences of YFP probability were expressed by continuous colormap for the subtracted YFP points. Vectorized YFP points and the subtracted YFP points were mapped onto the 3D whole brain shell using vedo. To visualize YFP^+^ cell distributions, the locations of YFP^+^ cells were first binned by 100 µm^3^ cubes in every brain sample, and then the averaged YFP^+^ cell density of every 100 µm^3^ cube was calculated across brain samples of the same experimental group. Every binned and averaged YFP^+^ cell cube (100 µm^3^) within particular regions was converted into a vectorized sphere using Numpy, Pandas and vedo. Each vectorized sphere expressed the information of the brain region to which the sphere belonged (discontinuous color map) and the size represented the normalized YFP^+^ cell density within the cube. Differential YFP^+^ cell density was determined by subtracting the YFP^+^ cell density of each vectorized point in the same location between the averaged mutant and WT brains. Scaled differences of YFP^+^ cell density were expressed by continuous colormap for the subtracted YFP^+^ cell spheres. Vectorized YFP cell spheres and the subtracted YFP cell spheres were mapped onto the 3D whole brain shell using vedo.

### Prediction of differential circuits established by Thy1-YFP projection neurons

Based on the correlation between YFP signals and YFP^+^ cell numbers in a given region, we screened the regions presenting differential innervation of YFP axons (see below for details). Then, we utilized the results of slice-based quantification in combination with 3D mapping and the spatial search service of the Allen Mouse Brain Connectivity Atlas ^22,23^ to predict the differential Thy1-YFP circuits in *Tbr1*^+/–^, *Nf1*^+/–^ and *Vcp*^+/R95G^ mouse brains compared to their respective WT littermates based on two assumptions. First, we based our prediction of Thy1-YFP circuits on the Allen Mouse Brain Connectivity Atlas, which constitutes a large number of axon tracing experiments in WT (C56BL/6J), Cre drivers and Ai75 (RCL-nT) reporter mice. It largely represents the mesoscopic connectome of WT mice. We hypothesized that the differential Thy1-YFP circuits of mutant mice simply contain different amounts of projecting axons from the original axon-projecting regions to the normal target regions and do not include misprojected axons. Second, since the source regions that project axons in the Allen Mouse Brain Connectivity Atlas are predominantly present in the right hemisphere, it influences the quantification of axonal projections originating in the left hemisphere. To get the maximum predictions of possible circuits, we hypothesized that hemispheric symmetry existed in the circuits we analyzed. In addition to the spatial search (Allen Mouse Brain Connectivity Atlas, see below) of a target region with the original location, we also performed a spatial search of the mirrored location at the other hemisphere (**Supplemental Figure 8**). The results of the two spatial searches were combined to obtain all possible ipsilateral and contralateral axon-projecting regions for a given target region in either the left or right hemisphere. Detailed methods are described in the following sections.

#### Step1: Identification of the putative target regions receiving differential YFP axon projections (also see **Supplemental Figure 6**)

For each brain region (excluding fiber tracts) in the compared group (mutant vs. WT), the binned (100 µm) mean YFP signal and mean YFP^+^ cell distributions along the AP axis of both mutant and WT samples were subjected to correlation analysis (using the scikit-learn Python package). If the mean changes in YFP signal (pixel number) along the AP axis were significantly correlated (significance level *P* < 0.05) with the mean changes in YFP^+^ cell number for both mutant and WT samples, Thy1-YFP cell number may thus reflect YFP signal abundance. Such brain regions were not considered candidate targets receiving differential axon projections of Thy1-YFP neurons. However, for a given region, if the changes in YFP signal along the AP axis were not significantly correlated (*P* > 0.05) with changes in YFP^+^ cell number in both mutant and WT mice, and the region also exhibited differential YFP signal distributions in the compared group, the region was deemed a putative target receiving differential YFP axonal projections. We identified all putative target regions in the compared groups (*Tbr1*^+/–^ vs. *Tbr1*^+/+^, *Nf1*^+/–^ vs. *Nf1*^+/+^, *Vcp*^+/R95G^ vs. *Vcp*^+/+^) to predict the upstream axon-projecting regions in the next step. Those regions are indicated in green font in **Figure 6** and summarized **in Supplemental Figure 7**.

#### Step2: Prediction of differential Thy1-YFP circuits based on the repertoire of putative target regions (also see **Supplemental Figure 8**)

We used the spatial search facility of the Allen Mouse Brain Connectivity Atlas (https://connectivity.brain-map.org) to predict the upstream axon-projecting regions. This system allows a user to select a seed location within the CCFv3 space to search all upstream regions with axonal projections to that seed location. To determine the location within the differentially-innervated regions for spatial searching, we used 3D mapping of YFP signal distributions. In both WT and mutant groups, the original YFP probabilities of 10-µm^3^ cubes in each 100-µm^3^ cube were first summed, i.e., binned into 100 µm^3^ cubes. For each 100-µm^3^ cube at the same location, the difference of summed YFP probability (mutant group – WT group) was calculated. Then, we identified the location of the 100-µm^3^ cube with the maximum difference and selected the cubes with the same differential tendency (mutant > WT: +1; mutant < WT: –1) as the sum of ΔYFP obtained in slice-based quantification. The location identified with the maximum difference and its mirrored location were then used for a spatial search in the Allen Mouse Brain Connectivity Atlas. The results were combined to obtain all possible ipsilateral and contralateral upstream regions. The putative upstream regions were further filtered by two criteria: (1) putative upstream regions having significantly different YFP^+^ cell distributions (sum of ΔYFP^+^ cell number) between WT and mutant groups; and (2) the summed ΔYFP^+^ cell number of the putative upstream regions having the same differential tendency as the summed ΔYFP signals of their axon-targeting region. After filtering, all qualified target regions in the two hemispheres and their corresponding upstream regions shown in the Allen Mouse Brain Connectivity Atlas were used to draw the differential Thy1-YFP circuits. For each putative target region, there are multiple axon tracing experiments containing the same upstream region but having different virus tracing locations. Therefore, we averaged the projecting densities of all the axon tracing experiments showing the same upstream– target region pairs and used the averaged projecting density to represent the connection strength. Finally, we defined the projecting difference between each upstream–target region pair by the following formula:

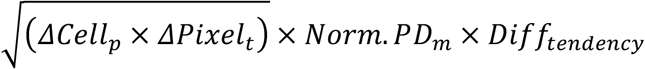

ΔCellp: Sum of ΔYFP^+^ cell number at the upstream projecting region;

ΔPixelt: Sum of ΔYFP pixel number at the target region;

Norm. PDm: Normalized mean projecting density of the upstream–target region pair (normalized to the maximum projecting density among all upstream to same target region pairs);

Difftendency: Differential tendency of summed ΔYFP signal at the target region (+1 or –1).

We used the information on all putative upstream axon-projecting regions, putative target regions, and the projecting differences to plot the differential Thy1-YFP circuits in both brain hemispheres in the pyCirclize Python package (**Figure 6**).

### Retrograde tracing of VP-projecting cells

#### Step1: Surgery for retrograde tracing

The mice (*Tbr1*^+/–^ and *Tbr1*^+/+^, 3-months-old) were deeply anesthetized via i.p. administration of a 5 mL/kg anesthetic mixture (Zoletil, 4 mg/mL; Xylazine, 2 mg/mL) and mounted on a stereotactic frame (Stoelting, Wood Dale, IL USA). After sanitization, followed by skull exposure, the bone and dura above the injection site (right VP) were removed and kept moist using cold 0.9% NaCl. A glass pipette (internal Ø 40 µm) loaded with 0.1% CTB-488 (Invitrogen, #C22841) was slowly inserted (10 µm/s) into the right VP [anterio-posterior axis (AP): –1.7 mm, medio-lateral axis (ML): 1.85 mm, dorso-ventral axis (DV): 3.75 mm] and one pressure pulse (10 psi, 30 ms duration) controlled by a pneumatic drug ejection system (PDES-02TX, NPI Electronic) was given to eject the retrograde tracer. After 10-min rest, the glass pipette was withdrawn slowly (10 µm/s). The mice underwent one week of recovery from surgery after appropriate suturing.

#### Step2: Acquisition of whole brain images and registration to CCFv3

One week after surgery, mice were deeply anesthetized via i.p. administration of 2,2,2-Tribromoethanol (0.7-0.8 mL per mouse), then transcardially perfused with PBS, followed by 4% PFA in PBS. The brains were collected and post-fixed overnight at 4 °C. After two days of dehydration in 30% sucrose in PBS, the brains were cryosectioned (coronal sections of 60 µm thickness) and kept in PBS. Correctly-injected samples were collected for further staining and imaging. Brain slices were washed in TBS three times (10 min each time) and the nuclei were labeled by DAPI staining (1 μg/mL). Serial slices of the entire brain were mounted in RapiClear CS (SunJin Lab, RCCS002) and imaged using a high-content imaging system (Molecular Devices) with a 4× objective lens. The exposure time for imaging the injection site and retrograde signals was 50 ms and 200 ms, respectively. Subsequent procedures, including image processing, registration to CCFv3, and auto/manual ROI corrections, are as described in detail above for Thy1-YFP images.

#### Step3: Comparison of injection sites (also see **Supplemental Figure 5B, 5C**)

To confirm that the injection sites were comparable between *Tbr1*^+/+^ and *Tbr1*^+/–^ mice, the K-Nearest Neighbors (KNN) classifier with 5-fold cross-validation was first used to evaluate the appropriate neighbors (k). For each proper k, the significance level was decided by comparing the accuracy between the original and shuffled data (1000 random permutations for each genotype).

#### Step4: Normalization of retrograde signals (also see **Supplemental Figure 5D**)

To extract the retrograde cell number, retrograde signals from image series with a long exposure time (200 ms) were first identified by pixel classification using ilastik (1.3.3post3). The retrograde signals were labeled and each pixel’s probability was returned via the Random Forest classifier. Subsequently, the object classification function in ilastik was used to classify the retrograde versus background signals. A group of pixels with the same instance segmentation were recognized as an object. The objects of the retrograde signals were labeled and the results were outputted as binary images. To count retrograde cell numbers, the spot detection function in Imaris (9.8.0) was used to detect and calculate the number of signals from binary images. The non-specific signals were removed manually.

There were three steps in obtaining the size of the area of injection signals. First, the injection site signals in short exposure time images (50 ms) were extracted via pixel classification, followed by object classification, and the binary mask of each slice was exported. Second, the pixel size of each object was counted, with the knee point on the cumulative curve of pixel size being detected using the Python package kneed, which acted as the threshold to remove most of the non-specific signals (i.e., objects with smaller pixel sizes). Third, a spatial limitation was set to select the pixel size of injection sites. The longest consecutive signals along brain slices and a rectangular area containing the VP region (2840.64 µm × 1971.2 µm in CCFv3 space) were set to extract the injection site signals.

For normalization, the retrograde cell number in each brain region was divided by the sum of pixel size in the injection site signals.

### Olfactory discrimination and preference tests

These behavioral tests were performed as described previously ^28^. Before the actual experimental test, two glass plates with plain filter papers were placed at the two ends of a home cage for 2 days for object habituation. In the discrimination test, six consecutive trials were conducted to assess olfactory sensation (trial 1), habitual adaptation (trials 2–5), and odor discrimination (trial 6). In the first five trials, 100 µM limonene (Cat. No. 8.1840, Merck) and mineral oil were separately spotted onto the filter papers placed at the two ends of the home cage for 5 min at 15-min intervals. In the sixth trial, the filter paper with mineral oil was replaced by filter paper spotted with 100 µM 2-heptanol (Cat. No. 8.20619, Merck). One week after the discrimination test, limonene and 2-heptanol were spotted individually onto filter papers, which were placed at the two ends of the home cage for 5 min in the preference test. Mouse behaviors were recorded using a camera installed on the lid of the cage. The total time spent sniffing each filter paper was measured to indicate the olfactory response to each odorant.

### Quantification and statistical analyses

Sample size n represents the number of brain samples from individual animals or the number of animals in the behavior assays for each experimental group. We did not use statistical methods to predetermine sample sizes, but our sample sizes are similar to those of previously published histology-based brain connectome ^24,50^ and behavioral ^28^ studies. Slice-based quantifications of YFP signal and YFP^+^ cell number distributions, and concomitant prediction of differential Thy1-YFP circuits, are based on n = 6 for each genotype of compared groups (three compared groups of mutants vs. WT littermates: *Tbr1*^+/–^ vs. *Tbr1*^+/+^, *Nf1*^+/–^ vs. *Nf1*^+/+^, *Vcp*^+/R95G^ vs. *Vcp*^+/+^). Retrograde tracing of VP-projecting cells was based on n = 7 for *Tbr1*^+/+^ and n = 7 for *Tbr1*^+/–^. Behavioral assays were based on n = 11 for *Nf1*^+/+^, n = 13 for *Nf1*^+/–^, n = 11 for *Vcp*^+/+^ and n = 11 for *Vcp*^+/R95G^. Exact n values for experimental groups in each analysis are presented in the figure legends. For data distributions, we did not use statistical methods to test data normality for the region-wise comparisons of YFP signal, YFP^+^ cell numbers and retrograde CTB spot distributions, but we assumed most of the quantified data is not normally distributed. Therefore, non-parametric statistical tests were used. For the behavioral assays, paired t-tests were used to analyze the olfactory discrimination data. Details of statistical tests are presented in the method sections describing each analysis. Data representing statistical parameters such as mean ± SEM are described in the figure legends. The test statistics (e.g., exact P values, Friedman chi-square statistics, and degrees of freedom) of region-wise comparisons (YFP signals, YFP^+^ cell numbers and retrograde CTB spot distributions) can be found in **Supplemental Table 1-7**, and the test statistics of behavioral analyses are given in **Supplemental Table 8**.

